# Discovering differential genome sequence activity with interpretable and efficient deep learning

**DOI:** 10.1101/2021.02.26.433073

**Authors:** Jennifer Hammelman, David K. Gifford

**Author notes:** **Corresponding Author** Correspondence to David Gifford.

## Abstract

Discovering sequence features that differentially direct cells to alternate fates is key to understanding both cellular development and the consequences of disease related mutations. We introduce Expected Pattern Effect and Differential Expected Pattern Effect, two black-box methods that can interpret genome regulatory sequences for cell type-specific or condition specific patterns. We show that these methods identify relevant transcription factor motifs and spacings that are predictive of cell state-specific chromatin accessibility. Finally, we integrate these methods into framework that is readily accessible to non-experts and available for download as a binary or installed via PyPI or bioconda at https://cgs.csail.mit.edu/deepaccess-package/.

**Author Summary:** Within the genome are the instructions to build all the cell types that make up the human body. However, understanding these instructions and how and when these instructions go wrong in cancer or genetically inherited disease is an open problem. Deep neural networks provide powerful models to learn the relationship between DNA sequence and functional consequence across many different cell types, such as whether a particular stretch of DNA is accessible and genes in that region can be expressed or is inaccessible and therefore genes are inactive. Despite these advances, a major setback in deep learning is that it is challenging to understand what patterns of DNA sequences a deep learning model has learned to associate with a particular genomic function, whether these patterns are significant, and how to determine whether these patterns are specific to a particular cell type or are general “housekeeping” patterns that function across many cell types. We introduce Expected Pattern Effect and Differential Expected Pattern Effect, two methods which allow us to evaluate the significance of particular patterns of DNA sequence features on models trained to predict function across multiple cell types, and apply this to problems of transcription factor binding and DNA accessibility across multiple cell types.

## Introduction

Neural networks have been increasingly helpful in understanding and predicting cellular function from genomic sequence in tasks such as transcription factor binding (1,2), chromatin accessibility (3,4), and histone modification (5). Previously, we showed that DeepAccess, an ensemble of neural networks trained to predict chromatin accessibility, is able to identify *de novo* DNA sequence motifs that influence local cell type-specific accessibility in the context of a massively parallel reporter assay (6). Here, we introduce an approach called the Expected Pattern Effect, which can be used to predict the effect of DNA sequence patterns such as known transcription factor motifs, combinations of motifs, or transcription factor spacing on the chromatin state of specific cell states. We also introduce a method to compare Expected Pattern Effects (EPE) between two conditions or cell states, which we call the Differential Expected Pattern Effect (DEPE). While the scientific community has yet to come to a consensus on a precise definition of cell state or cell type, for the purposes of this work we define cell type as a shared characteristic gene expression profile. For example, motor neuron cells are defined by characteristic expression of the motor neuron and pancreas homeobox gene Hb9 (7). A cell state is similarly defined through gene expression response, but refers to a shared change in gene expression as a response to external stimuli. We focus on cell type in this work, but the methods we introduce are also readily applicable to analysis of cell state.

## Results

Recovering interpretable biological hypothesizes from trained deep neural network models has been a challenge given the complexity of the models. Contemporary approaches for interpreting deep learning models of genomic sequence features utilize three main approaches (Fig 1A):

1. Interpretation of model convolutional filters as position weight matrices (3,4,8,9).
2. Interpretation of model and individual sequences for important nucleotides using model saliency and motif generation from input sequences (10–13) or saturating in silico mutagenesis (2,5,12).
3. Model response to in silico “designed” sequences (3,14–17).

**Fig 1.**
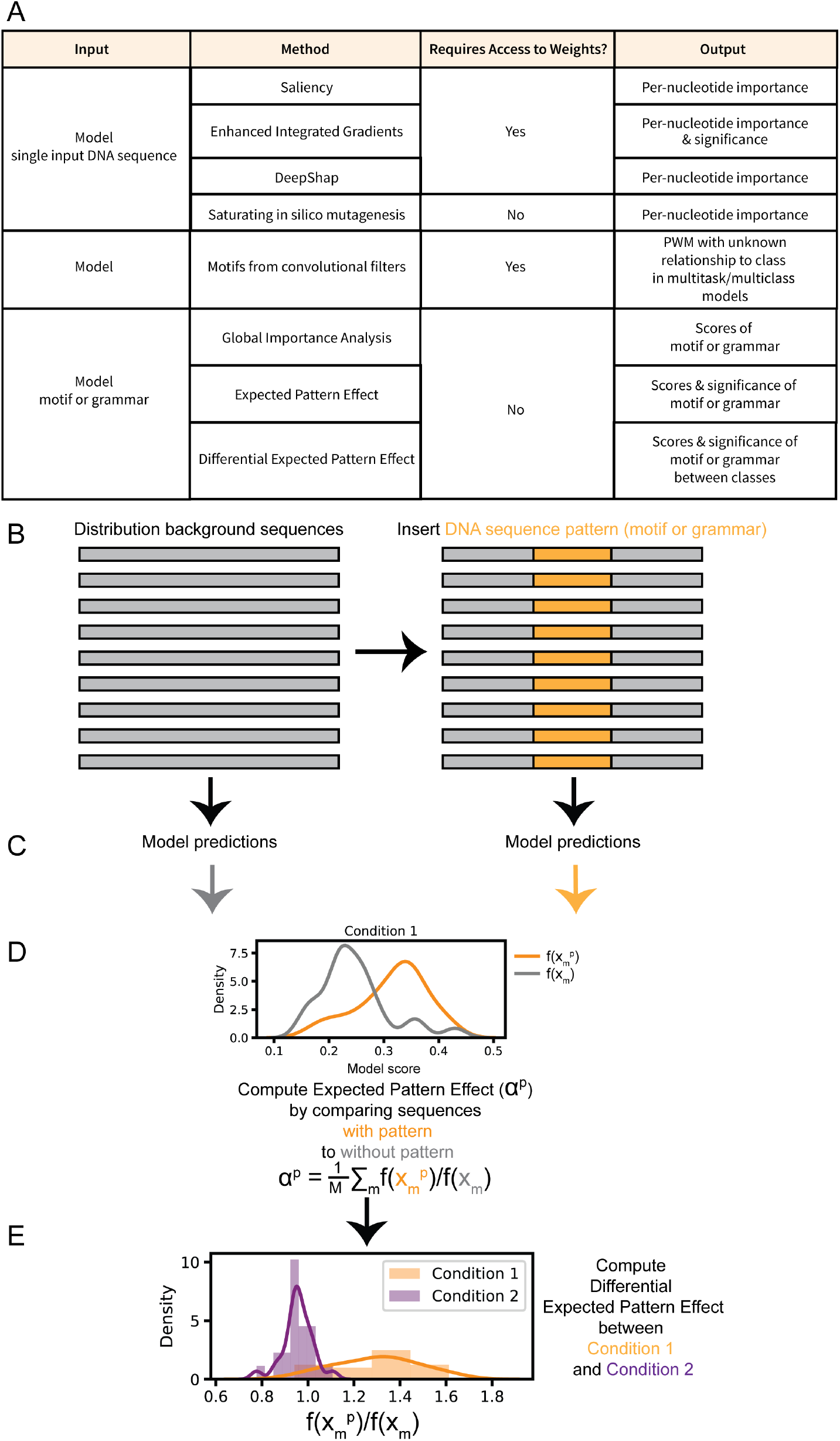
Expected Pattern Effect and Differential Expected Pattern Effect determine the significance of feature patterns such as transcription factor motifs or transcription factor grammar for multitask models. (A) Comparison of interpretation methods shows 3 categories of interpretation methods for deep learning models based on input. Some methods operate on an individual sequence input, model pair, others turn learned convolutional filters into DNA sequence motif position weight matrices and then must assign class relevance post-hoc, or methods Global Importance Analysis and Expected Pattern Effect score the overall class importance of DNA sequence patterns such as transcription factor motifs or grammar such as combinations or spacing of transcription factor motifs. (B) Computation of Expected Pattern Effect and Differential Expected Pattern Effect starts by selection of a sequences that represent a “background” from the natural distribution of sequences. A pattern of interest either a transcription factor or combination or spacing of transcription factors is artificially inserted into each background sequence. (C) Model predicts functional signal for “background” sequences and sequences with inserted transcription factor motif. In our case we use a DeepAccess model, but principles for computing EPE and DEPE can be applied to any multitask classification model. (D) Average ratio between scores of background sequences with pattern of interest (in orange) and without pattern of interest - i.e. background sequences (in grey) determines the Expected Pattern Effect. (E) Differential Expected Pattern Effect can be computed by comparing Expected Pattern Effect between different classes - i.e. conditions or cell types.

Methods in class 1 can generate novel potentially important motifs from the trained weights of convolutional filters, but filters have been known to represent partial motifs (9) and additional analysis is required to interpret the significance of these motifs on cell type-specific activity. Methods in class 2 such as DeepSHAP (10,18) and Enhanced Integrated Gradients (13) provide means to identify DNA sequence features associated with a single condition and differentially between two conditions (Fig 1A), but these methods function to identify DNA sequences features within an individual input sequence, so the importance of these sequence features is dependent on the surrounding nucleotides within in the input DNA sequence and do not represent the overall expected effect of a particular pattern such as a transcription factor motif on cell type-specific accessibility. In the case of models operating on DNA sequence, we have knowledge *a priori* that particular combinations of features (patterns) that could have important effects, such as transcription factor motifs. Interpretation methods make use of our *a priori* knowledge about the patterns of features such as DNA motifs and relationships between patterns such as spacing to investigate what features the model has learned are overall most important through evaluating model response to “designed” DNA sequences *in silico*. However, these approaches have yet to explore how to assign the significance of these effects and how to interpret whether these effects are differential between two conditions, such as cell types, when applied to multi-task classification models.

Here we present a method of assigning both effect scores and significance values to DNA sequence feature patterns of interest, such as transcription factor motifs or grammars such as particular transcription factor motifs and motif spacings, that differentiate between cell types or conditions from a multi-task neural network (Fig 1B). Significant differential sequence features can suggest hypotheses that relate specific DNA binding factors to cell state. Thus, these factors can be candidates for follow up studies on their effectiveness in directed differentiation protocols designed to produce specific cells types.

In contrast to methods that analyze a single input sequence or a model, we build upon the use of “designed sequences” to create a computational method to measure the learned differential effect of a transcription factor motif or other DNA sequence pattern on the output of a DeepAccess model (Fig 1A). These patterns can include a combination of transcription factor motifs. Many deep learning models are multi-task models that simultaneously predict more than one function. For example, they may simultaneously predict the binding of multiple transcription factors or the chromatin accessibility of multiple cell types. The output of these models is often a probability. Thus, for a binary measurement, *y*, over *N* training examples, we can estimate the overall expected probability of *y* to be:

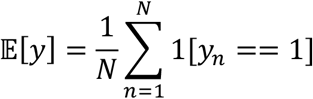

Since we intend to compare the effect of genomic sequence features with imbalanced class data, we have to account for the relative effect of average class prediction probability. Thus, we take into account the average class prediction probability in our formalization of the Expected Pattern Effect, α, of a genomic pattern, *p*, such as a transcription factor motif (Fig 1B). Given a model that predicts function from sequence 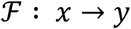, where *x* is a sequence of nucleotides and *y* represents a corresponding binary functional measurement, the effect of pattern, *p*, can be estimated by the impact of the insertion of *p* into the sequence *x* to make the new sequence *x^p^*, both of which are within the data distribution *D*, and is given by the ratio of expectations:

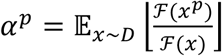

While EPE is a general method that can be applied to any multi-task classification model, here we apply it to investigate cell type-specific chromatin accessibility using a multi-task DeepAccess model *f_c_* : *x* → *y_c_*, to approximate 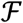, were *y_c_* denotes the model predictions on functional task, *c*. DeepAccess can be trained and thus predict on one or more classes *c*. We select *M* sequences to approximate *D* and estimate 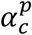 according to:

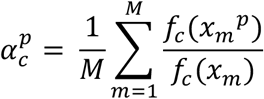

We define the significance of the Expected Pattern Effect (EPE) as the signed-rank test statistic of the ratios:

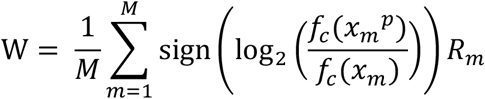

where *R_m_* represents the rank of the absolute log ratio of sequences with the pattern insertion,

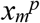, to sequences without the pattern insertion, *x_m_*.

We then compute the Differential Expected Pattern Effect between two classes *c_1_* and *c_2_* based upon the average log_2_ ratio of predictions:

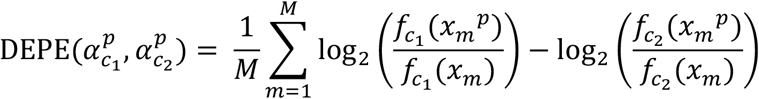

In addition to computing the magnitude of the DEPE between two classes, we compute the significance of this effect:

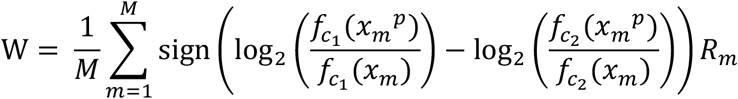

where *R_m_* represents the rank of the absolute log ratio of the expected pattern effect of the pattern, *p*, differentially between two classes *c_1_* and *c_2_*, on a sequence *m*.

Our goal is to apply the DEPE to discover cell type-specific transcription factor activity from chromatin accessibility, but to first validate that DEPE can correctly identify differential transcription factor activity, we apply the DeepAccess architecture to predict DNA binding from ChIP-seq data. In contrast to chromatin accessibility where multiple or specific combinations of transcription factors may be responsible for activity, ChIP-seq should produce sequences enriched by the binding of a single DNA binding protein. Therefore, if a DeepAccess model has successfully learned differential DNA sequence patterns between two ChIP-seq experiments, DEPE should identify the DNA binding motifs of those transcription factors.

We evaluated the effectiveness of the DEPE by evaluating how well it could identify sequence motifs that were responsible for differential ChIP-seq binding. We trained a single DeepAccess model to predict the DNA sequences bound by six ChIP-seq experiments for the transcription factors ELF1, MEF2A, CTCF, USF1, TCF12, and NRF1 in mouse erythroleukemia (MEL) cells from ENCODE (19). The trained DeepAccess model takes as input a DNA sequence and outputs the predicted binding of six transcription factors to the input sequence (six outputs). The DeepAccess model was trained with the 10,000 most significantly bound genomic regions from GPS (20,21) binding event calls. Each region is classified as being bound by either neither, one, or multiple transcription factors based on whether it overlaps a GPS binding event. After successfully training the DeepAccess model, we ran our method for DEPE to extract differential patterns which result in significant differential binding predictions, using the HOCOMOCOv11 (22) database of transcription motifs as our patterns. The transcription factor motifs matching the transcription factor ChIP are found to have significant DEPE (Fig 2), except in the case of differential motif discovery between NRF1 and TCF12, where the TCF12 motif was not discovered as a differential pattern between NRF1 and TCF12 binding (Fig 2C). The DMRT1 and PO5F1 motifs had the strongest DEPE on TCF12 binding relative to NRF1, suggesting that TCF12 may be binding indirectly through partnering with other proteins, including a transcription factor recognizing ACAAT DNA motif which is shared by DMRT1 and PO5F1 in MEL cells. Despite the large overlap in binding sites between NRF1 and TCF12, DEPE is able to identify the NRF1 motif as a differential pattern between NRF1 and TCF12 binding, indicating that our approach is able to identify differential motifs even when two classes share a majority of genomic regions (Fig 2C; Fig S1). The number of motifs that are identified as significantly differential is asymmetric for comparisons between ELF1 and MEF2A (Fig 2A) and USF1 and CTCF (Fig 2B). This can be explained by the large size of the bHLH-ZIP (USF1) and Ets (ELF1) transcription factor families, each encompassing greater than 20 transcription factors with highly similar binding motifs within the HOCOMOCOv11 transcription factor database (22). However, the same is not true for NRF1 which also has asymmetric differential transcription factor motifs when DEPE is computed between NRF1 binding and TCF12 binding. It appears that NRF1 binding sites may frequently contain binding motifs for other transcription factors such as Klf/Sp factors, and Gata which may indicate cooperative binding relationship. This in combination with the large number overlap of Nrf1 and Tcf12 binding sites suggests Nrf1 and Tcf12 binding may be regulated by logic beyond the presence or absence of their DNA binding motifs. We also show that under class imbalanced data, EPE and DEPE generate robust estimates of transcription factor effects in contrast to Global Importance Analysis (15), which takes a related approach but uses a difference instead of a ratio to estimate the effect of a pattern (Fig S2).

**Fig 2.**
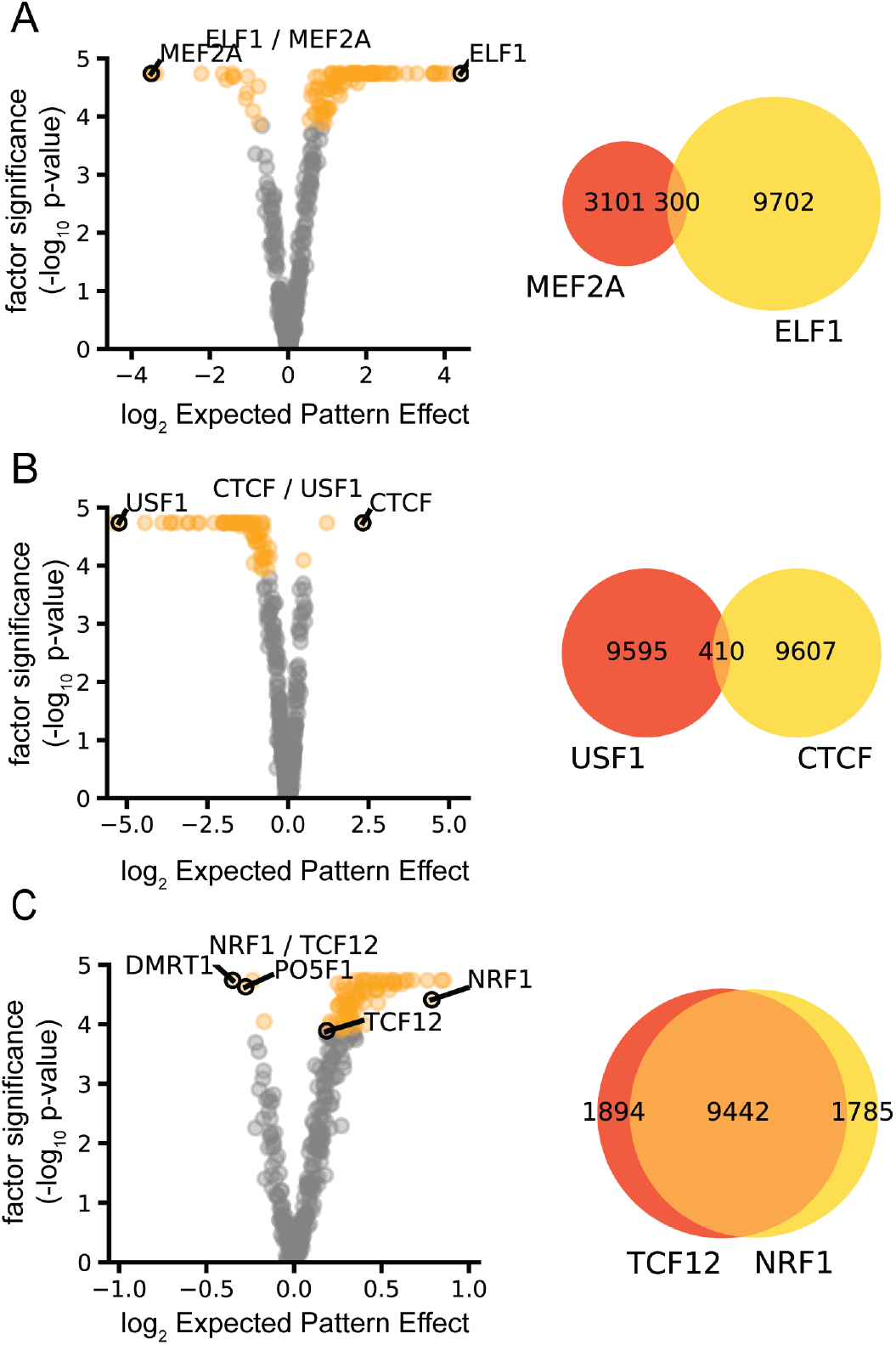
Differential Expected Pattern Effect identifies condition-specific transcription factor activity from DeepAccess models. DeepAccess trained on mouse ChIP-seq data identifies correct motifs driving differential transcription factor activity for (A) ELF1 and MEF2A, (B) CTCF and USF1, (C) NRF1 and TCF12. In the scatter plot each point represents one of 356 HOCOMOCOv11 transcription factor motifs. Scatter plot y-axis is unadjusted rank ratio significance, x-axis is the Differential Expected Pattern Effect. Orange points are significant transcription factor motifs (*p* < 0.05) under Bonferroni multiple hypothesis correction. Venn diagrams show number and overlap of ChIP-seq binding events used as training and testing data for DeepAccess.

We then used EPE and Differential EPE to investigate chromatin accessibility. First, we hypothesized that we can use EPE to investigate preferential spacing of Oct4 and Sox2 motifs within accessible DNA in stem cells, as these two transcription factors are known to cooperatively bind. We tested the EPEs for multiple spacings of Oct4 and Sox2 motifs (Fig 3A) and find the DeepAccess model predicts the highest Expected Pattern Effect from the spacing of Pou5f1 and Sox2 that is consistent with the spacing between the DNA motifs of Pou5f1 and Sox2 co-binding events from ChIP-seq binding data (Fig 3B). We then examined Differential EPE for all transcription factor motifs in the HOCOMOCOv11 database between stem cells and motor neurons (Fig 3C) and find homeobox transcription factors such as Hoxc9, Meis1, and Hoxa9 are differentially important for motor neuron accessibility consistent with the important role that Hox genes play in motor neuron development (23,24), whereas transcription factor motifs for Maz and Sp5 are differentially important in stem cells, with Sp5 and Maz sharing a similar DNA binding motif and Sp5 playing an established role downstream of the WNT pathway to maintain self-renewal in stem cells (25,26). We found that EPE and DEPEs for transcription factors is generally not affected by the position of insertion within the designed sequences, except when the site of insertion is at the very beginning of the DNA sequence (Fig S3). Thus, we used the center of the DNA sequence as our insertion site for all subsequent experiments. Additionally, we examined how selection of background sequences to approximate D affects EPE and DEPE, and found that EPEs are smaller when distribution-approximating sequences are selected from accessible or inaccessible parts of the genome (Fig S4) but are more or less invariant to specific annotations (i.e. promoters) compared to randomly selected DNA. Since we are most interested in motifs driving accessibility, we use randomly sampled closed DNA as our distribution sequences. We also examined Differential EPE between liver and fibroblast, identifying transcription factor motifs Hnf1a, FoxD3, and Hnf1B as being differentially important to liver (Fig 3D), consistent with the role of hepatocyte nuclear factors (Hnfs), including Fox, in liver differentiation (27–29). Finally, we compute the Differential EPE for each of the ten cell types relative to the other nine cell types, for all 356 mouse core HOCOMOCOv11 transcription factor motifs (Fig 3E), and find that Differential EPE of transcription factors identifies transcription factors that are known lineage regulators, such as LIM homeobox transcription factors in dopaminergic neurons [DN] (30,31) and motor neurons [MN] (32–34), as well as Hox transcription factors which are specific to spinal motor neurons. JUN/FOS/AP-1 family members c-Jun, c-Fos, and Fosl1are likely to play a role in fibroblast [FIBRO] identity as knockdown of these transcription factors facilitated reprogramming of fibroblast to stem cells (35) and over-expression led to reduced reprogramming efficiency to stem cell state and the maintenance of accessibility of fibroblast-specific enhancers (36). Additionally, we find Fox transcription factors in both pancreatic [aPN, bPN] and liver cells [LIV] which is consistent with the role of Fox in pan endoderm differentiation and liver and pancreatic cell differentiation (28,37–39). Finally, hierarchical clustering using the Differential EPE for significant transcription factors showed relationships between cell types such as alpha pancreatic and beta pancreatic cells, and motor neuron and dopaminergic neurons which share more similar Differential EPEs (Fig 3E).

**Fig 3.**
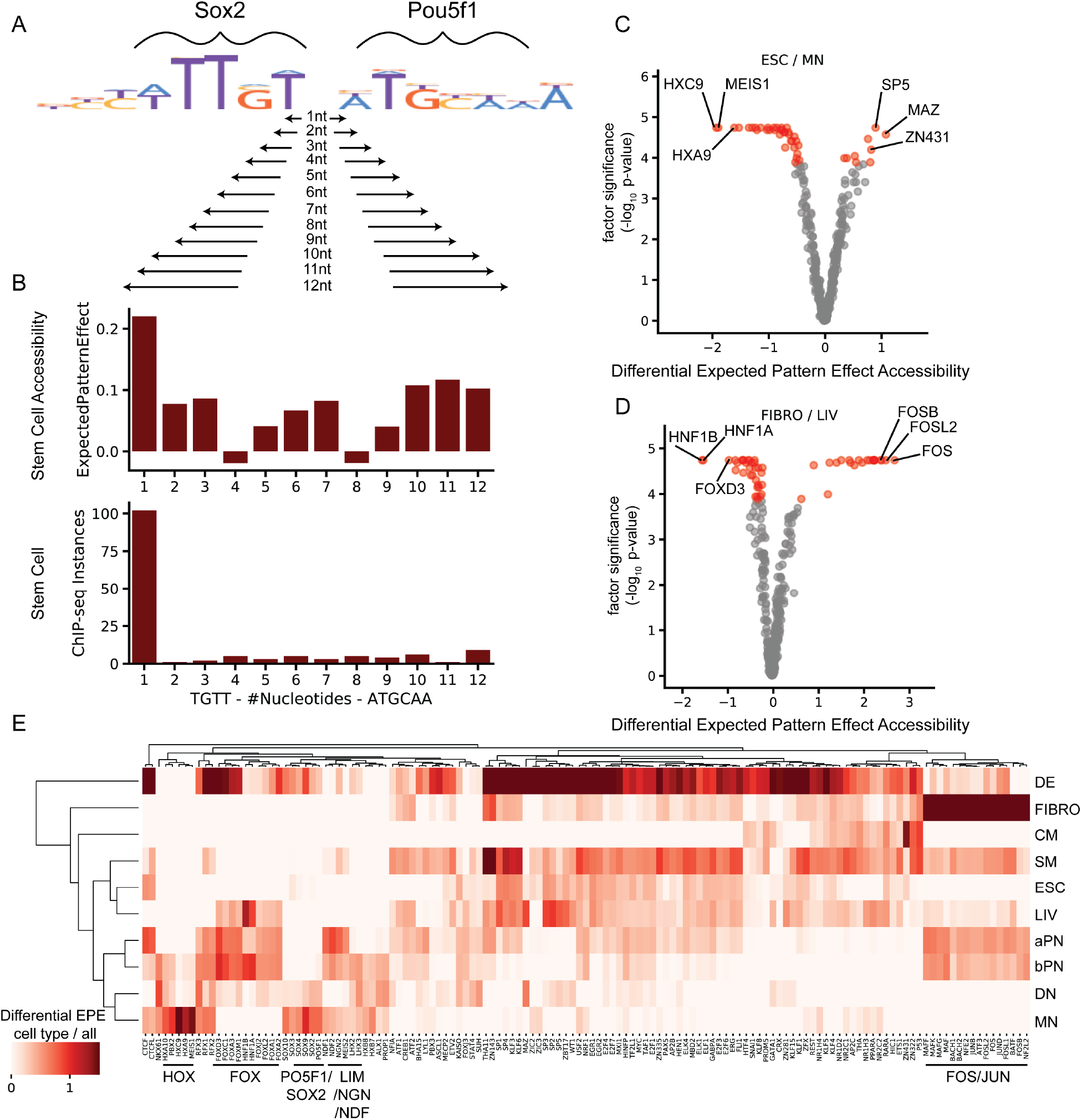
Expected Pattern Effect and Differential Expected Pattern Effect identify preferential spacing and cell type-specific transcription factor activity from DeepAccess models trained on chromatin accessibility data from ten cell types. (A) Testing all possible spacing between Sox2 transcription factor and Pou5f1 (Oct4) transcription factor motifs input as patterns to compute Expected Pattern Effects from a DeepAccess model. (B) DeepAccess Expected Pattern Effect of Sox2 and Pou5f1spacing on predicted stem cell accessibility (top) is consistent with preferential spacing of Sox2 and Pou5f1 motifs from ChIP-seq binding data of Sox2 and Pou5f1 co-binding events (C) Differential EPE of transcription factor motifs between stem cells (ESC) and motor neurons (MN) identifies known MN-specific transcription factors such as Hox TFs and Meis. (D) Differential EPE of transcription factor motifs between fibroblasts (FIBRO) and liver cells (LIV) identifies liver-determining factors such as Hnf1 and Fox transcription factors. Scatter plot y-axis is unadjusted rank ratio significance, x-axis is the Differential Expected Pattern Effect (log_2_ fold change in Expected Pattern Effect between cell types), with each point representing one of 356 HOCOMOCOv11 transcription factor motifs. Red points are significant (*p* < 0.05) transcription factor motifs under Bonferroni multiple hypothesis correction. (E) Transcription factors showing significant cell type-specific EPEs for each of the ten cell types relative to all other cell types. Heatmap intensity indicates the log_2_ fold change in Expected Pattern Effect between each cell type and the average of all other cell types.

We have increased the utility of our method by providing an executable version of DeepAccess for the Linux operating system as well as a python package available via PyPI and bioconda (Fig 4A). Using Keras (v2.4.3) with a Tensorflow (v2.4) backend for neural network deployment, the DeepAccess package automatically parallelizes to take advantage of multiple cores on a CPU or GPU. The DeepAccess system takes as input multiple bed files (Fig 4B), each labeled by experimental condition (i.e., cell type, condition, genomic signal), or fasta files with labels for each fasta input sequence (Fig 4C). The DeepAccess system then splits the data into training and test data, performs training with early stopping criteria based on a held-out validation set of the training data (Fig 4D). Performance metrics and predictions are reported for training and test data (Fig 4E). Finally, we include our methods for interpretation with testing EPE and Differential EPE between conditions (Fig 4F) and differential importance of individual nucleotides (Fig 4G). Additionally, the DeepAccess system is also able to make predictions on new input DNA sequences (Fig 4H).

**Fig 4.**
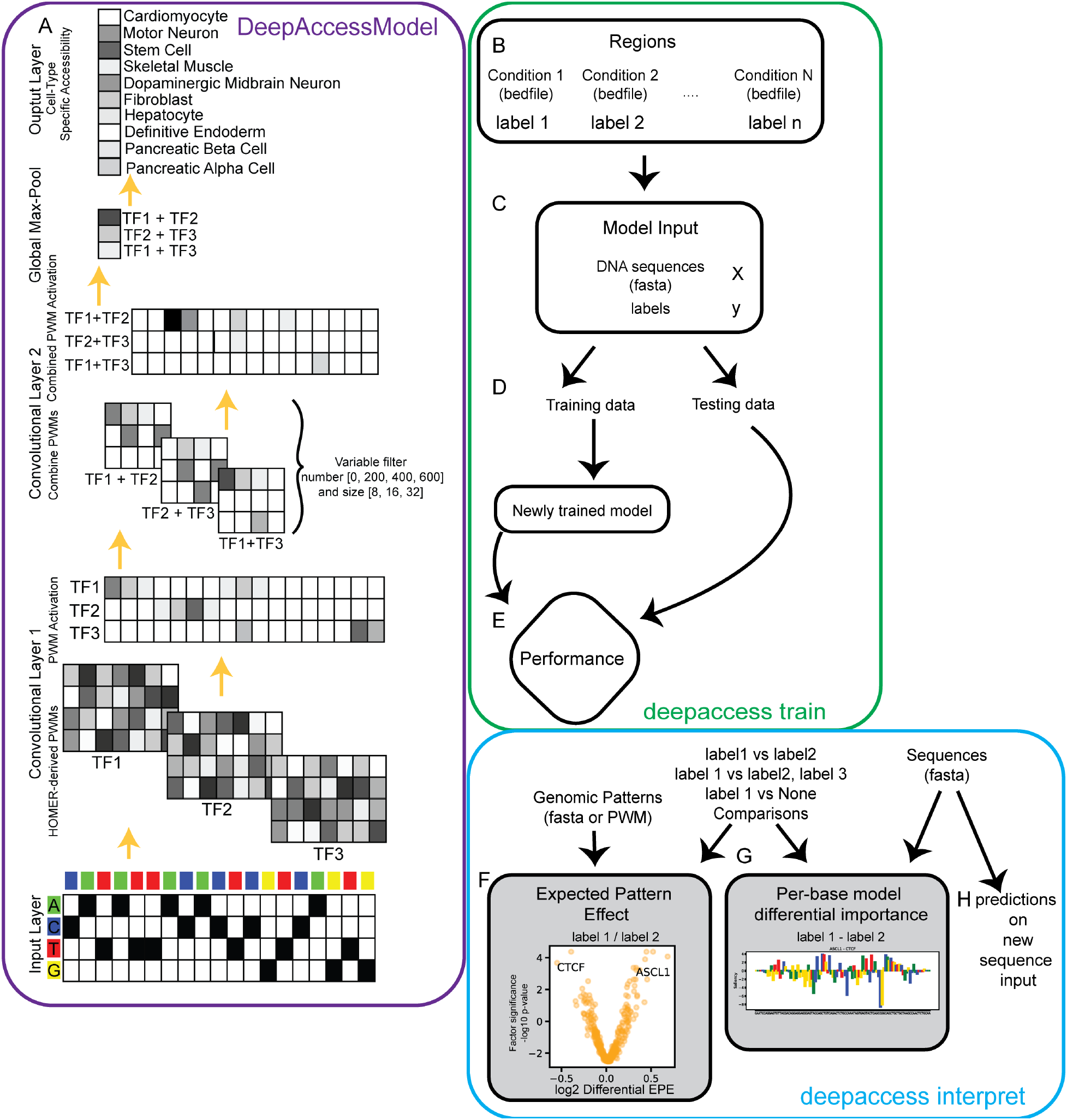
Accessible and interpretable deep learning with DeepAccess. (A) Core DeepAccess model example trained to predict accessibility from 100nt DNA sequence using ATAC-seq data from 10 mouse cell types. Input layer is one-hot encoding of DNA sequence. Convolutional layers scan for transcription factor motifs using position weight matrices derived from the HOMER database. Ensemble of models with variable filter number and size in the second convolutional layer combine transcription factor motifs. Global max pooling layer condenses results into maximally activating position for convolutional filter. Output is the multi-task accessibility which can be interpreted as the probability that an input DNA sequence will be accessible in each cell type, state, or condition. The DeepAccess system takes as input either (B) genomic regions from multiple conditions with labels, or (C) DNA sequences as fasta files and labels. (D) Data is split into training and testing, and model is trained with early stopping criteria based on a validation subset of ATAC-seq data. (E) Model performance is computed. Finally, results of newly trained model can be interpreted using (F) ExpectedPatternEffect and DifferentialExpectedPatternEffect to evaluate activity of genomic patterns for a given condition or between conditions, (G) per-base model differential importance scores between conditions and (H) predictions on new input sequences.

The DeepAccess system, including methods for estimating EPE and Differential EPE, is available as a PyPI package and on bioconda. We also provide a binary of the DeepAccess system which avoids typical installation problems with the evolution of Python packages by packaging all dependencies including the Python interpreter within our software. Even when using package managers such as pip and conda which are present with frameworks such as Selene (40), Kipoi (41), and pyyster (42), problems with incompatibility can still arise with typical package managers as a consequence of python, package manager, and operating system updates.

## Discussion

While previous work has explored the contributions of neural network architecture, training methodology, and model interpretation to learn the relationship between DNA sequence and differential DNA binding (12), these neural networks were trained to only predict cell type specific binding. In contrast, DEPE can be used to interpret DNA sequence features influencing differential accessibility in cases where models have been trained to predict accessibility across tens, hundreds, or thousands of unique cell types. In these cases, explicitly training models on each differential comparison between cell types would be computational expensive and time intensive. We also note that Expected Pattern Effect resembles other methods for extracting pattern effects from deep neural networks (3,15), except that we compute Expected Pattern Effect to permit the comparison of pattern effects between conditions, which is important for identification of cell type-specific or condition-specific sequence features and show that our use of ratio to compare effects is more robust to analysis of cell type-specific transcription factor activity under class imbalance.

While DeepAccess was developed specifically for identifying cell type-specific sequence features from chromatin accessibility, Differential Expected Pattern Effect can be used to discover condition-specific sequence features from many types of experimental genome-wide sequencing data. We expect that DeepAccess will provide a useful approach for discovering sequence features that drive differential cell fates, and that the system we provide will make this capability accessible to a wide audience.

## Methods

### ChIP-seq / ChIP-nexus read processing and binding event calling

Reads were trimmed for adaptors and low-quality positions using Trimgalore (Cutadapt v0.6.2) (43) and aligned to the mouse genome (mm10) with bwa mem (v0.7.1.7) (44) with default parameters. Duplicates were removed with samtools (v1.7.2) (45) markdup, and ChIP binding events were called with GPS (v3.4) (20,21) with default parameters.

### DeepAccess ChIP-seq training

We used as input to DeepAccess 200bp regions centered at the GPS-called ChIP-seq binding event. DeepAccess then uses these regions to create a training data set of 87,114 100bp sequences which can be labeled as binding events for any combination of the 6 tested transcription factors, including regions where none of the 6 transcription factors are predicted to bind. We define a region as a ChIP binding event for a transcription factor if more that 50% of the 100bp region overlaps the 200bp region centered at a GPS binding event. The DeepAccess model was then trained for 3 epochs with default parameters.

### DeepAccess Model trained on ATAC-seq for 10 mouse cell types

Methods for DeepAccess and differential nucleotide importance with DeepAccess were previously described in Hammelman et al., 2020. Briefly, we trained DeepAccess on ATAC-seq data from ten cell types: stem cell, fibroblast, hepatocyte, endoderm, beta pancreatic cell, alpha pancreatic cell, cardiomyocyte, skeletal muscle, dopaminergic midbrain neuron, and spinal motor neuron using binary cross-entropy loss (multi-task classification) with 4,220,507 genomic regions for training: 3,220,507 regions were open in at least 1 cell type, and 1,000,000 regions were closed in all cell types (randomly sampled from the genome). Chromosome 18 and chromosome 19 were held out for validation and testing, respectively. To define training regions for DeepAccess, we generate 100bp genomic windows across the entire mouse genome. We define a region as accessible in a given cell type if more that 50% of the 100bp region overlaps a MACS2 (46) accessible region from that cell type. The pretrained DeepAccess model is available as a Zenodo record at https://zenodo.org/record/4908895#.YL6YpR0pDfY.

For ATAC-seq processing, reads were trimmed for adaptors and low-quality positions using Trimgalore (Cutadapt v0.6.2) (43). Reads were aligned to the mouse genome (mm10) with bwa mem (v0.7.1.7) (44) with default parameters. Duplicates were removed with samtools (v1.7.2) markdup (45) and properly paired mapped reads were filtered. Accessible regions were called using MACS2 (v2.2.7.1) (46) with the parameters -f BAMPE -g mm -p 0.01 --shift −36 --extsize 73 --nomodel --keep-dup all --call-summits. Accessible regions that overlapped Encode blacklist regions ENCSR535HHO were excluded from downstream analysis.

### Preferential spacing of Oct4 and Sox2

ChIP-nexus binding data for Oct4, Sox2, and patch cap control (17)(17) were downloaded as fastqs from the Nucleotide Read Archive (see Table S1 for accession numbers). ChIP-seq processing was performed as described (see ChIP-seq / ChIP-nexus read processing and binding event calling). After, all GPS events where Sox2 or Oct4 binding sites were less than 50nt were considered as close co-binding events. using bedtools (v2.29.2) (53) intersect. Events were scanned for the first instances of TTGT (Sox2 DNA sequence motif) and ATGCAA (Oct4 DNA sequence motif), and the distance between the last nucleotide of the Sox2 DNA sequence motif and the first nucleotide of the Oct4 DNA sequence motif was defined as our distance between TF sites.

### Selection of Synthetic Distribution of Sequences

For all experiments we used as our synthetic distribution sequences 24 randomly selected genomic locations that were not overlapping ATAC-seq accessible regions in all ten mouse cell types. We provide these regions with the DeepAccess executable as default backgrounds for new experiments.

## DeepAccess Executable

DeepAccess executable was generated with pyinstaller (v4.1).

## Data availability

No new data were produced by this study. Previously published ChIP-seq and ATAC-seq data are reported in Table S1 and Table S2 of methods.

## Code availability

Binaries for DeepAccess training and interpretation with ExpectedPatternEffect and differential saliency is available http://cgs.csail.mit.edu/deepaccess-package/. Source code is available at https://github.com/gifford-lab/deepaccess-package

## Acknowledgements

We thank members of the Gifford lab and Wichterle lab for helpful discussions and contributing data. We gratefully acknowledge funding from 1RO1HG008363 (D.K.G.), 1R01HG008754 (D.K.G.), 1R01NS109217 (D.K.G.), and National Science Foundation Graduate Research Fellowship (1122374) (J.H.).

## Contributions

Conceptualization, J.H. and D.K.G.; Methodology, J.H.; Software, J.H.; Formal Analysis, J.H. and D.K.G.; Investigation, J.H.; Resources, D.K.G.; Data Curation, J.H.; Writing - Original Draft, J.H.; Writing - Review & Editing, J.H., and D.K.G.; Visualization, J.H.; Supervision, D.K.G.; Funding Acquisition, J.H. and D.K.G.

## Ethics Declarations

The authors declare no competing interests. The software described here is available under a MIT license and is free for non-commercial and commercial use.

## Supplemental Legends

Table S1. ChIP-seq data used to test DeepAccess Differential Expected Pattern Effects and for preferential spacing of Oct4 / Sox2 in stem cell accessible DNA.

Table S2. ATAC-seq data used for pre-trained DeepAccess model.

Fig S1. All pairwise differential Expected Pattern Effects from ChIP-seq shows correct TF motifs are identified in most comparative analyses.

Fig S2. Stability of estimated motif effects with Global Importance Analysis (subtraction) and Expected Pattern Effect (ratio) shows EPE more accurate on multi-task data sets with class imbalance. Scatter plots of GIA (A-C) or EPE/DEPE (D-F) of effect sizes of 356 HOCOMOCOv11 mouse motifs on predictions of CTCF binding from a trained DeepAccess model on balanced class data (x-axis) compared to 2x, 10x, and 30x more CTCF binding sites : USF1 binding sites (y-axis) (A) GIA of CTCF binding (B) GIA of USF1 binding (C) GIA of CTCF vs USF1 differential binding (D) EPE of CTCF binding (E) EPE of USF1 binding (F) DEPE of CTCF vs USF1 binding.

Fig S3. Effect of motif insertion position on EPE. (A) EPE of Hoxc9 motif on EPE from DeepAccess model trained to predict motor neuron chromatin accessibility shows EPE of Hoxc9 motifs are low for positions in the beginning of the DNA sequence and high for positions at the middle and end of the DNA sequence. (B) Scatter plot of EPEs for 356 HOCOMOCOv11 mouse transcription factor motifs when insertion is at position 0 compared to position 50 indicate early insertion site in DNA sequence results in different EPE values for transcription factors, with some having lower EPEs and others having higher EPEs. (C) Scatter plot of EPEs for 356 HOCOMOCOv11 mouse transcription factor motifs when insertion is at position 70 compared to position 50 indicate early insertion site in DNA sequence results in similar EPE values for all transcription factors. (D) Heatmap of Pearson’s r of EPEs for DeepAccess motor neuron accessibility for insertions at all starting positions within the DNA sequence shows most insertion positions result in highly similar EPEs except for early insertion positions. (E) Heatmap of Pearson’s r of DEPEs for DeepAccess motor neuron vs fibroblast differential accessibility for insertions at all starting positions within the DNA sequence shows most insertion positions result in highly similar EPEs except for early insertion positions. (F) Row z-score normalized EPEs for DeepAccess motor neuron accessibility at each position show most motifs have higher EPEs when inserted at a position in the center of the DNA sequence. Some motifs (like Sp motifs) have higher EPEs at the beginning of the DNA sequence.

Fig S4. Choice of M sequences approximating sequence distribution affects EPE scores. (A) Estimates of EPE for 356 HOCOMOCOv11 transcription factor motifs on predicted motor neuron accessibility from a learned DeepAccess model. M sequences are selected randomly from regions closed in all 10 cell types, open in all 10 cell types, promoters which are closed in all 10 cell types, and promoters which are open in all 10 cell types. Shade indicates distribution size (M=10, M=25, M=100). Boxplot box indicates median and quartiles and whiskers extend to 1.5 times the inter-quartile range. Dots are outliers outside 1.5 times the inter-quartile range. (B) Estimates of significance of EPE for size and type of sequence used to estimate total sequence distribution. (C) Comparison of EPEs for 356 HOCOMOCOv11 transcription factor motifs for default (randomly selected 24 closed) compared to randomly selected closed (row 1), randomly selected open (row 2), promoter closed (row 3), and promoter open (row 4) shows while values of EPE dependent on selection of sequence distribution, rank remains relatively invariant.

