## Supplemental Figures and Tables for "Discovering differential genome sequence activity with interpretable and efficient deep learning"

| Protein (ChIP-seq) | Data Source | Identification |
| --- | --- | --- |
| MEF2A | ENCODE | ENCFF433FQJ; ENCFF423RCQ |
| ELF1 | ENCODE | ENCFF186NAS; ENCFF592ERV |
| USF1 | ENCODE | ENCFF996MWJ; ENCFF550MZG |
| TCF12 | ENCODE | ENCFF880XUQ; ENCFF432YTR |
| NRF1 | ENCODE | ENCFF845NFO; ENCFF205UPE |
| CTCF | ENCODE | ENCFF001NRQ; ENCFF001NRO |
| IGG | ENCODE | ENCFF001NCH; ENCFF001NCF |
| Oct4 | GSE137193 | GSM4072776 |
| Sox2 | GSE137193 | GSM4072777 |
| Patch cap | GSE137193 | GSM4072780 |

**Table S1. ChIP-seq data used to test DeepAccess Differential Expected Pattern Effects and for preferential spacing of Oct4 / Sox2 in stem cell accessible DNA.**

| Cell type (ATAC-seq) | Source | Identification |
| --- | --- | --- |
| Stem cell ATAC-seq | (De Dieuleveult et al. 2016) | GSE64825 |
| Fibroblast ATAC-seq | (Cheloufi et al. 2015) | GSE66534 |
| Skeletal Muscle ATAC-seq | (Ramachandran et al. 2019) | GSE123879 |
| Cardiomyocyte ATAC-seq | (Quaife-Ryan et al. 2017) | GSE95763 |
| Endoderm ATAC-seq | (Cernilogar et al. 2019) | GSM3223249 |
| Hepatocyte ATAC-seq | ENCODE | ENCSR609OHJ |
| Pancreatic alpha/beta cell ATAC-seq | (Lawlor et al. 2017) | GSE99954 |
| Dopaminergic Midbrain Neuron ATAC-seq | (McClymont et al. 2018) | GSE122450 |
| Spinal Motor Neuron ATAC-seq | Closser et al. 2021 in revision | GSE178466 |

**Table S2. ATAC-seq data used for pre-trained DeepAccess model.**

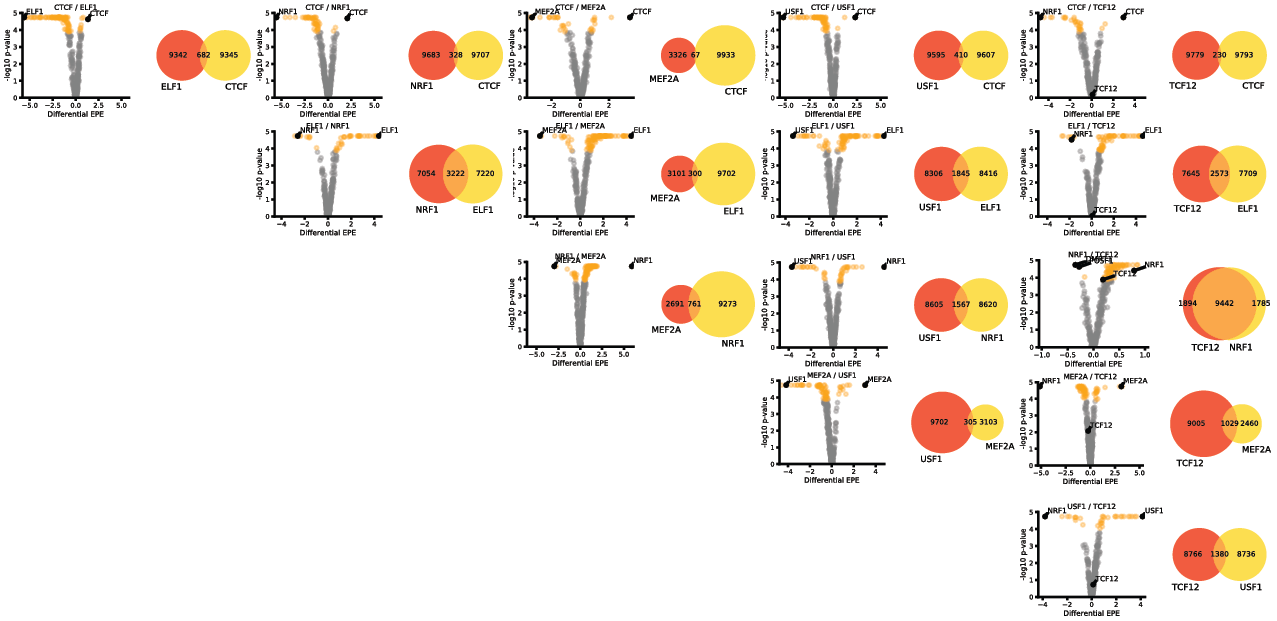

**Figure S1.**

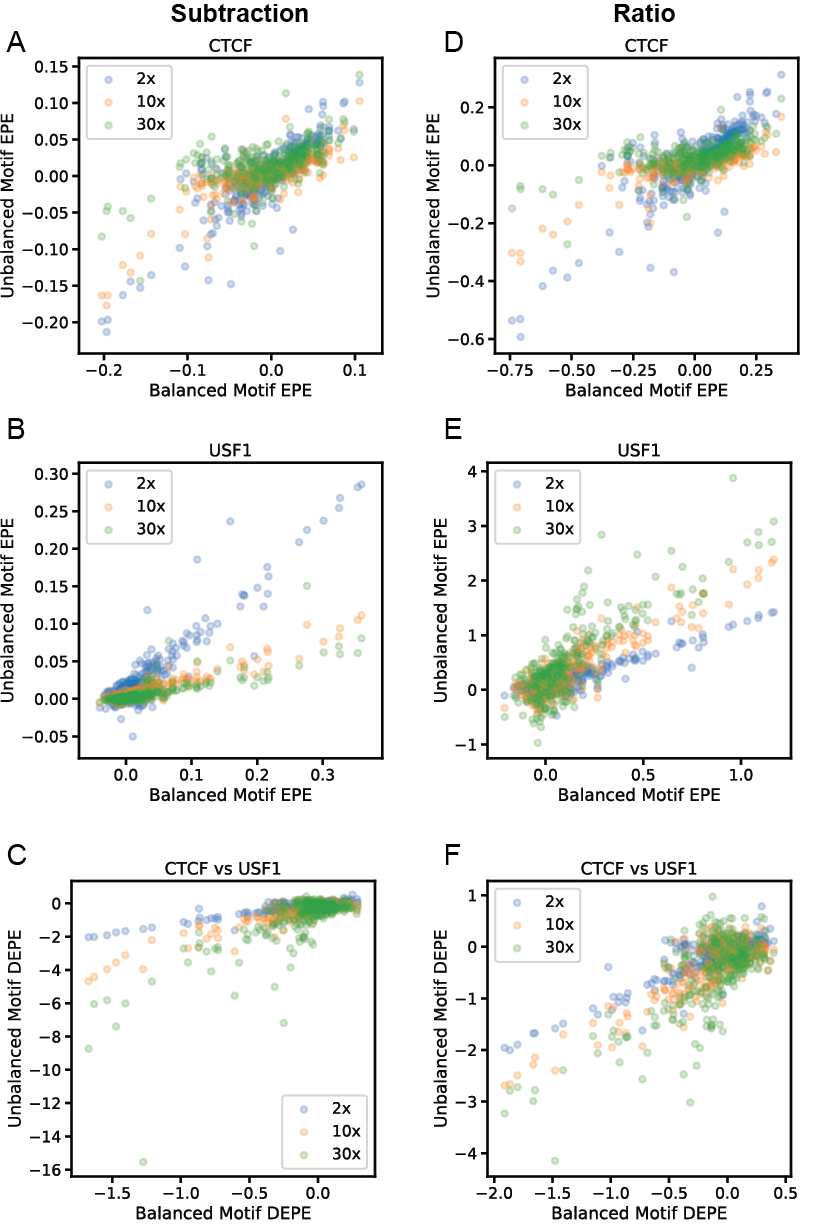

**Figure S2.**

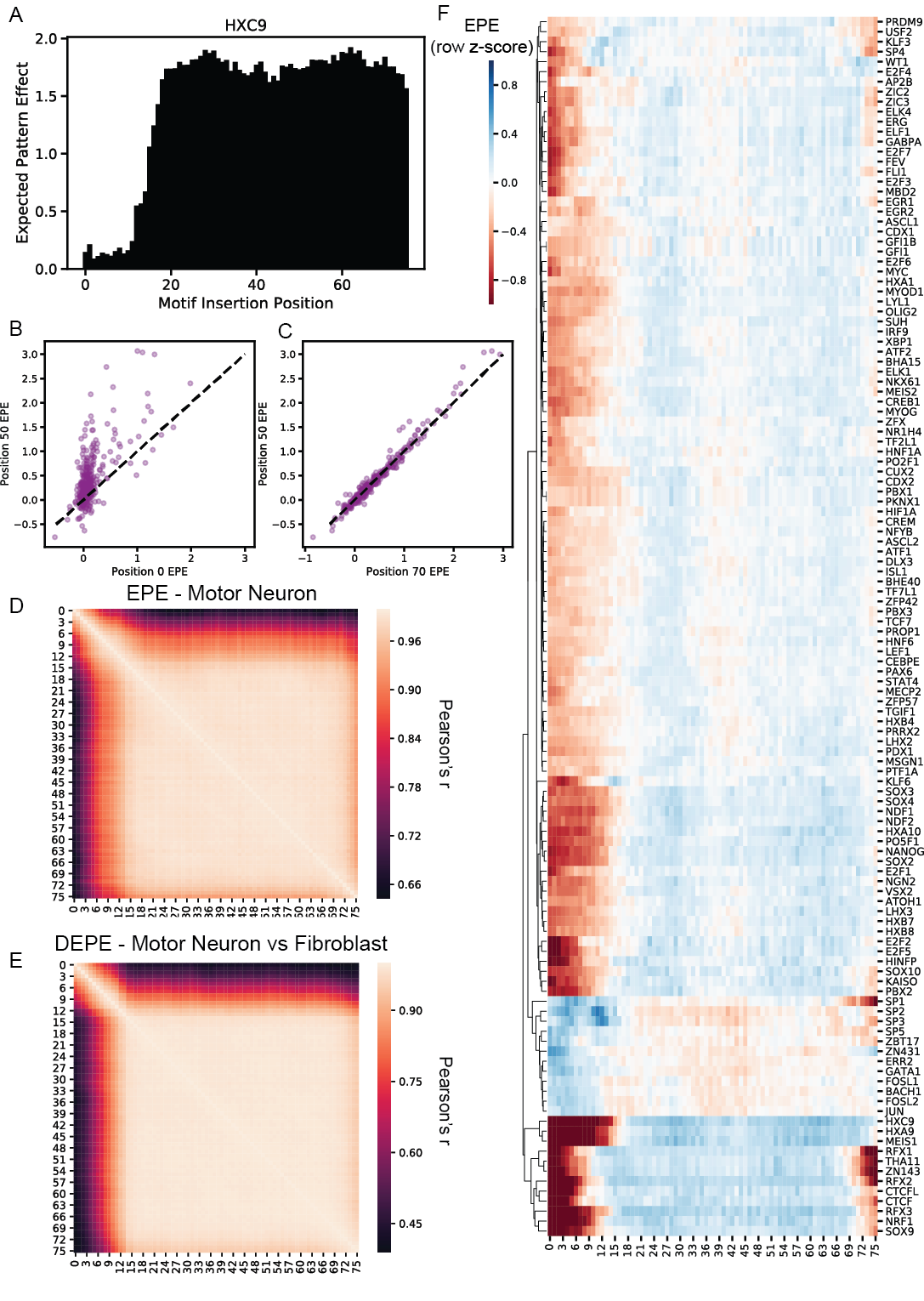

**Figure S3.**

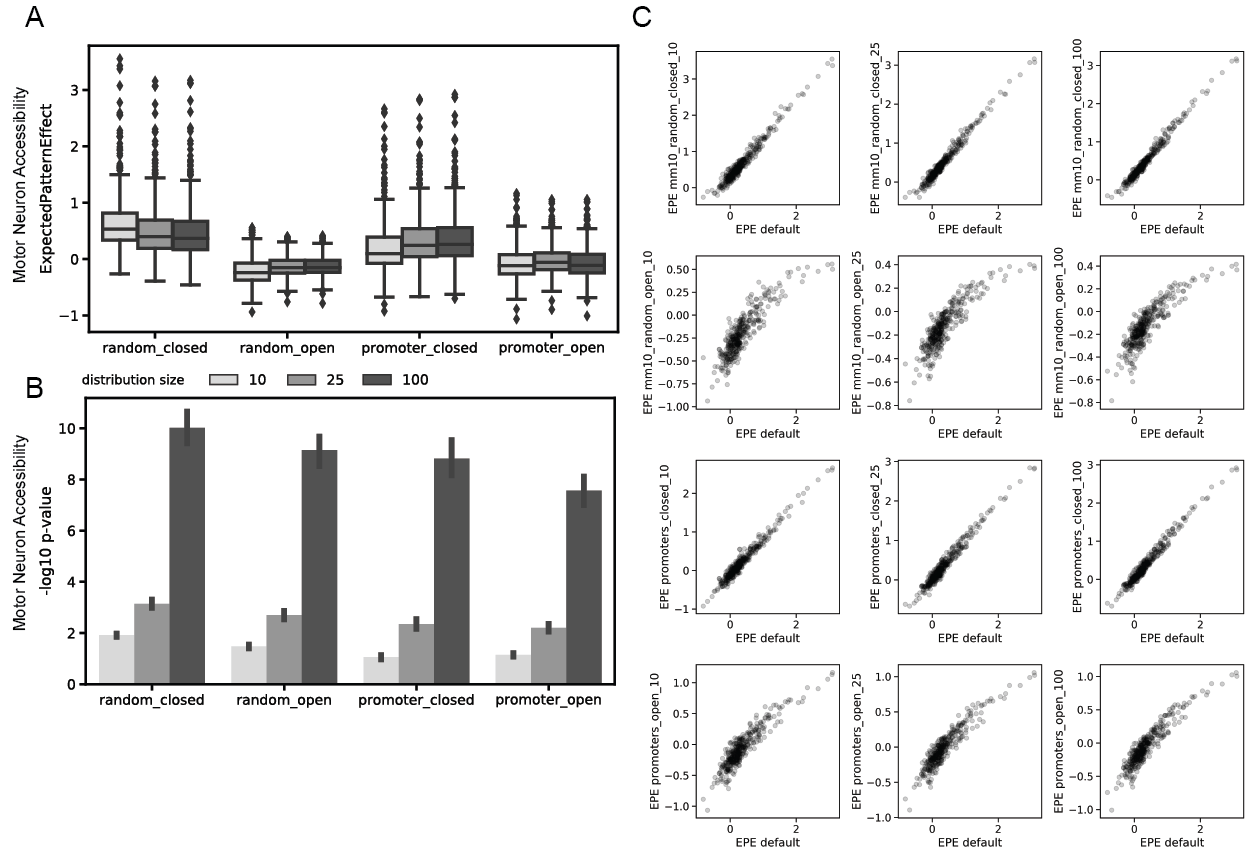

**Figure S4.**
